# The Impact of Control Interface on Features of Heart Rate Variability

**DOI:** 10.1101/2021.05.07.443181

**Authors:** Mahdieh Nejati Javaremi, Di Wu, Brenna Argall

**Affiliations:** Northwestern University, Shirley Ryan AbilityLab, Chicago, Illinois; Northwestern University, Chicago, Illinois

**Keywords:** Assistive Robotics, Heart rate variability, Workload Prediction

## Abstract

Shared human-robot control for assistive machines can improve the independence of individuals with motor impairments. Monitoring elevated levels of workload can enable the assistive autonomy to adjust the control-sharing in an assist-as-needed way, to achieve a balance between user fatigue, stress and independent control. In this work, we aim to investigate how heart-rate variability features can be utilized to monitor elevated levels of mental workload while operating a powered wheelchair, and how that utilization might vary under different control interfaces. To that end, we conducted a 22 person study with three commercial interfaces. Our results show that the validity and reliability of using the ultra-short-term heart-rate variability features as predictors for workload indeed are affected by the type of interface in use.

## 1 INTRODUCTION

Assistive machines such as powered-wheelchairs can increase the independence and quality of life for persons living with motor impairments. When a person is fitted for a powered wheelchair, the seating clinician will take their unique constraints and abilities into consideration when choosing the *control interface* they will use for operating their machine. The selected control interface can affect not only *how* the person operates their machine, but also what *challenges* they may face [1].

Incorporating robotics autonomy within assistive machines can improve *human autonomy* by collaborating with the human user to ensure safety and relieve physical and cognitive burden. The robotics autonomy operates by observing the external world and the user through sensors such as cameras and range sensors mounted on the assistive machine. Each person is distinct in their abilities and experiences and therefore requires unique amounts and types of assistance. Therefore, robotics autonomy should vary how and when it steps in to assist based on the observations made about the world and the user, and knowledge it has about the user’s individual preferences.

One measure which can be potentially used as an implicit cue for when the autonomy should step in is mental workload. Mental workload has been defined in multiple ways due to the diverse perspectives investigating this phenomenon, and subsequently, various measurement formalisms have been proposed [2]. Studies in various domains have shown that heart-rate variability (HRV) correlates with cognitive workload [3, 4]. However, the majority of this work studies tasks that last more than five minutes and often during rigorous physical activity, while many wheelchair driving tasks occur on much smaller time-scales and the only physical activity of the user is the operation of the control interface.

Using HRV as a measure of cognitive load requires the use of physiological sensors to continuously monitor the state of the user. Although wireless ECG sensors are now commercially available, there are still challenges with using these sensors to measure HRV during activities of daily living: (1) the sensors need to be placed at precise locations on the body, (2) the sensors typically only last a few hours of constant use and need to be recharged, (3) sweat and sensor displacement can negatively affect the accuracy of the sensor readings, and (4) conductive gel is typically required for improved sensor readings which if dried can also negatively affect the accuracy of measured signals.

The long term aim of this work is to design robotics autonomy that is able to discern changes in the user’s cognitive demand from features of their control input signal. Commercially available interfaces fall into three broad categories: joysticks, switches, or sip-n-puff straws. These machines can be operated in various ways—including hand motion, head tap, or respiration—which may result in different sensor readings from physiological sensors while navigating the same tasks. How this variation affects HRV workload calculations has yet to be explored and is an aim of this paper. Furthermore, the signals generated through these interfaces can have different characteristics in terms of proportionality, dimensionality, continuity of the signal, to name a few, which can affect how the user input signal may correlate with the HRV signals.

To accomplish our larger goal we first need to establish an online ground-truth measure of workload—as opposed to an episodic measures such as the NASA-TLX. As a first step towards our larger goal we provide the following contributions:

1. Identify whether ultra-short-term (UST) HRV features can accurately predict cognitive load during wheelchair navigation.
2. Identify whether UST HRV features are impacted by the type of control interface used.
3. Identify how the workload prediction accuracy of UST HRV features are affected by the spread and density signals as a direct result of interface selection.

## 2 BACKGROUND

In this section, we will summarise relevant prior work.

### 2.1 Cognitive Load

Various definitions of mental workload exist in the literature without a clear consensus. Tao et. al. [5] distinguishes between metal workload, physical workload, and task load. They assert that mental workload (MWL) conveys stresses resulting from task demands which reflect an individual’s subjective experience performing a task given external conditions and constraints. This is contrasted to the physical workload that result from stresses on the human’s physical body and task load that signifies the amount of work an individual must perform. In this work, we follow the definition put forward by Young and Stanton which suggested that MWL refers to “the level of attentional resources required to meet both objective and subjective performance criteria, which may be mediated by task demands, external support and past experience”[6].

A review article by Charles et. al [2] performed a meta analysis over all common physiological measures of workload. Their results found that there is no universal solution for measuring mental workload and no single stand-out method was recommended. However, they found that physiological sensors are able to capture the experience of the user during different tasks and contrasted this to subjective measures which can be influenced by how the individual perceived their own performance. Giannakakis et. al.[7] developed a stress recognition system using HRV that classified stressful and non-stressful scenarios with 84.4% accuracy under a variety of stressors, including under cognitive load while performing the Stroop Color Word Task. Ultra short-term HRV measures were also used to identify mental stress in students during an academic exam[3]. In another study, mental workload as measured by the visual analog scale of fatigue and the NASA TLX score was shown to alter HRV during an hour-long letter recognition task[8].

### 2.2 Heart-Rate Variability

HRV measures the variation in time intervals between adjacent heartbeats and is considered to be an index for neurocardiac function[9]. This variation arises from the complex dynamics between heart-brain interactions and non-linear autonomic nervous system processes. Interdependent regulatory mechanisms such as (but not limited to) autonomic balance, blood pressure, gas exchange, and vascular tone all contribute to HRV, but operate on different time scales, hence the need for different HRV measures. As these regulatory mechanisms are employed when we adapt to environmental and psychological challenges, HRV can help us to index an individual’s mental workload[10]. For instance, higher levels of resting vagally-mediated HRV are linked to performance of executive functions like attention and emotional processing by the prefrontal cortex[9].

A variety of HRV metrics exist to probe at different mechanisms that underlie HRV. Time domain metrics measure the time variation between consecutive heartbeats. Many metrics drop abnormal heartbeats, deriving metrics only from the normal to normal heartbeat intervals (NN intervals). Frequency domain metrics estimate the absolute or relative power in a certain frequency band. Non-linear and entropy metrics quantify the regularity and complexity of successive heartbeat intervals[10].

### 2.3 Workload Monitoring for Robotics Autonomy

A variety of assistive robotic wheelchairs have been proposed and shown to reduce the workload and cognitive burden of wheelchair drivers[11] using post-task subjective measures of workload. Although the knowledge that the assistance can reduce workload post-hoc is important, the assistance can be more beneficial if it can implicitly track the workload of the user online, and only step in when the workload exceeds an acceptable threshold. Towards this, Lamti et. al. extracted features from electroencephalogram sensor signals that model mental fatigue[12]. To our knowledge there is no study that has modeled workload using HRV features during powered wheelchair navigation with various interfaces.

## 3 MATERIALS AND METHODS

This section provides a detailed description of the research design and procedures used in the experiment.

### 3.1 Hardware

The study was conducted using a Permobil powered wheelchair equipped with an on-board computer, two RGB-D sensors, and wheel encoders. Three of the most common interfaces used for controlling powered wheelchairs were included in the study[13]. The selection used in this study were (1) an ASL 533 Compact joystick (ASL, TX, USA), (2) 105 electronic head array system (ASL, TX, USA), and (3) sip/puff switch (Origin Instruments, TX, USA) (Fig. 1). We used BioStamp RC sensors (MC10, MA, USA) to measure electrocardiogram (ECG) and accelerometer signals. The ECG sensor was attached in the LEAD I orientation as shown in Fig. 2 and signals were measured at a frequency of 250*Hz*.

**Figure 1:**
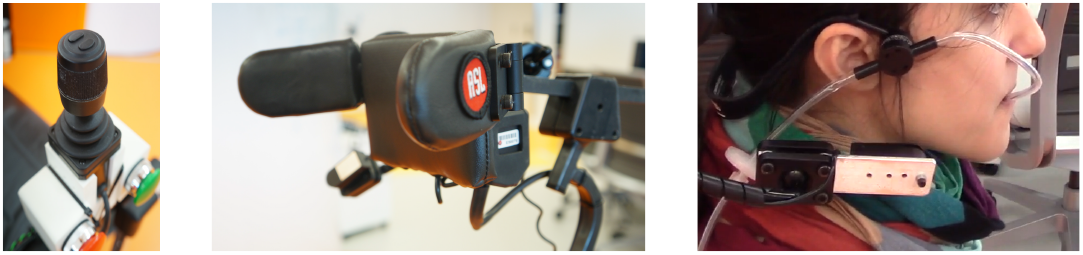
Control interfaces used in the study. *From left to right:* 3-axis joystick, head array device, sip/puff switch.

**Figure 2:**
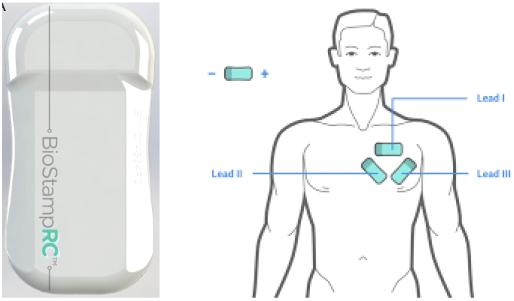
Wearable sensor used in this study. The BioStamp RC sensor can be configured to measure ECG, EEG, accelerometer and gyroscope signals. The sensor was placed in the Lead I configuration.

### 3.2 Participants

We recruited 23 participants: 9 with spinal cord injury (SCI) (41.6 ± 13.9 years, levels C3-C6, complete and incomplete) and 14 uninjured (31.6 ± 9.1). One of the uninjured subjects was excluded from the analysis due to excessive noise in the sensor signal. The experiment was approved by the Northwestern University Institutional Review Board (STU00207312) and informed consent was obtained from all participants. A questionnaire screened subjects for alcohol and coffee consumption, exercise prior to the study session and smoking habits.

### 3.3 Experimental Procedure

The study consisted of three sessions—one session per interface. The uninjured subjects took part in all three study sessions, while the SCI subjects only used the interfaces they were able to safely control. Each session began by measuring the baseline HRV of the participant at rest, followed by a training phase where participants were familiarized with the operation of the specific interface and the wheelchair dynamics. This was followed by the testing phase which involved wheelchair navigation through a track with seven tasks: two doorway traversals, ramp ascent and descent, avoiding a dynamic obstacle, and traversing a sidewalk with drop-off from wide and narrow ends (Fig 3). The order of the tasks within the circuit and the order of the interface sessions were randomly counter balanced across all subjects. At the end of each trial session, participants responded to a NASA-TLX questionnaire to measure subjective workload [14], and a post-task questionnaire to rank the relative difficulty of the tasks. The session concluded by again measuring resting HRV of the participant after completing the circuit.

**Figure 3:**
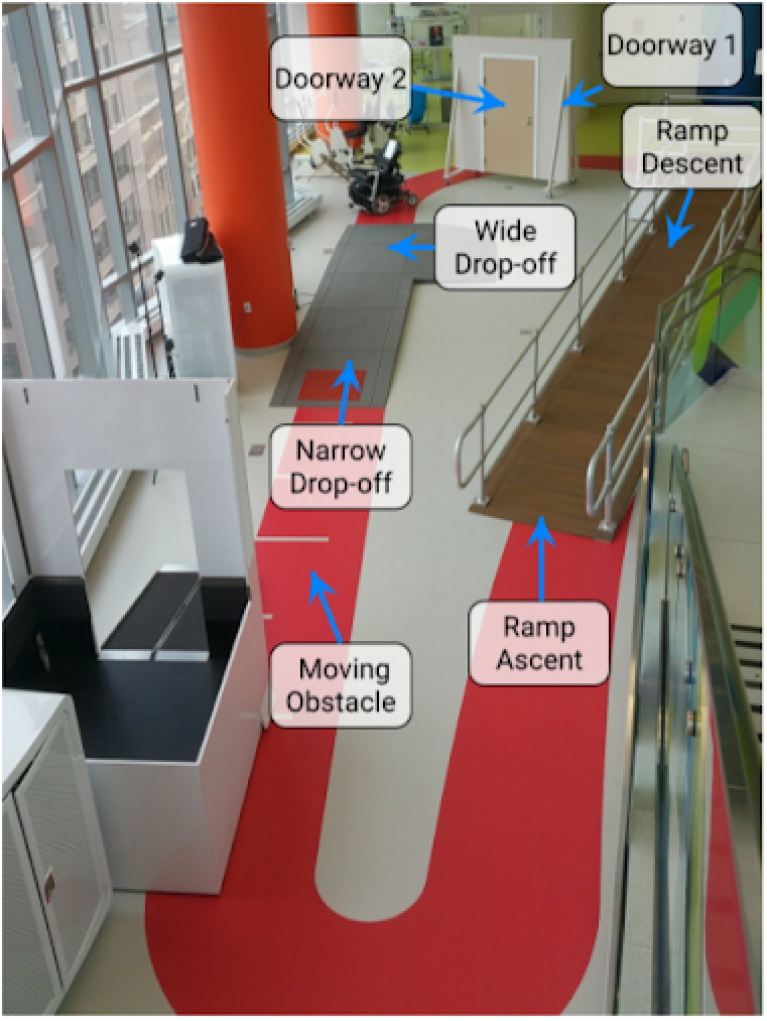
Wheelchair navigation track. Tasks include doorway traversal, ramp descent and ascent, drop-off avoidance, and dynamic obstacle avoidance.

### 3.4 HRV Metrics

The raw ECG data was processed using the Kubios HRV Premium analysis software. The processing steps included visual inspection of the signal and manual noise removal, correcting ectopic beats, outlier removal, and time series trend removal (Fig 4). The R-R intervals were automatically extracted from the raw ECG signals using the Kubio QRS detection function based on the Pan-Tompkins algorithm [15].

**Figure 4:**
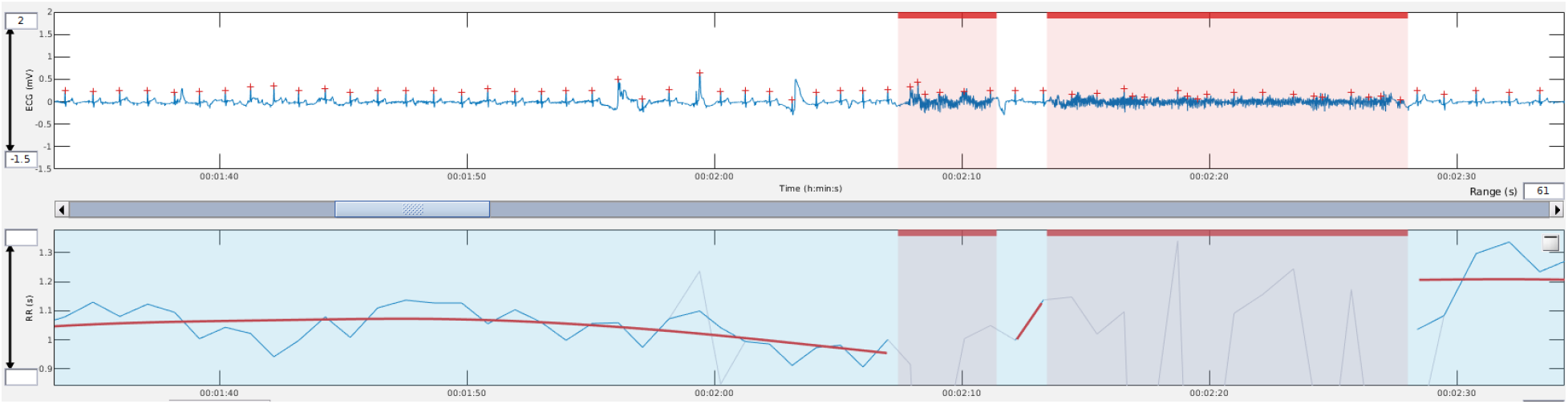
The raw ECG signal (top). Noisy segments were marked (red background) and removed from the analysis. R-R intervals were extracted from the QRS signal (bottom).

Due to the nature and duration of wheelchair tasks, we chose ultra-short-term (UST) (under 5 minutes) HRV features. HRV metrics were generated from the detected R-R intervals using the PhysionNet Cardiovascular Signal Toolbox[16]. These metrics were calculated using 30 and 60 seconds sliding windows at 1 second increments. Previous studies have investigated the reliability of various UST HRV features[17, 3, 18, 19, 20]. While these studies looked at individual feature correlations between short-term and ultra-short-term metrics, we are interested in whether these features can predict workload. The UST features used in our analysis are summarized in Table 1.

**Table 1:** Heart rate variability features.

| HRV Metric | Unit | Description |
| --- | --- | --- |
| **Time Domain Metrics** |  |  |
| NNmedian | ms | Median of all NN intervals |
| NNmode | ms | Mode of all NN intervals |
| NNvariance | ms | Variance of all NN intervals |
| NNmean | ms | Mean of all NN intervals |
| RMSSD | ms | Square root of the mean of the sum of differences between adjacent NN intervals |
| lnRMSSD | ms | Natural log of the square root of the mean of the sum of differences between adjacent NN intervals |
| SDNN | ms | Standard deviation of all NN intervals |
| **Frequency Domain Metrics** |  |  |
| hf | ms<sup>2</sup> | Power in high frequency range ($0.15 \leq hf < 0.4$ Hz) |
| lf | ms<sup>2</sup> | Power in low frequency range ($0.04 \text{ Hz} \leq lf < 0.15$ Hz) |
| lfhf | % | Ratio of lf to hf |
| **Entropy Metrics** |  |  |
| ApEn | a.u. | Approximate entropy, measure of regularity and complexity of NN time series |
| SampEn | a.u. | Eliminates ApEn bias toward regularity |
| **Non-linear Metrics** |  |  |
| SD1 | ms | Standard deviation of the projection of the Poincaré plot on the line perpendicular to the line of identity |
| SD2 | ms | Standard deviation of the projection of the Poincaré plot on the line of identity |
| SD1SD2 | % | Ratio of SD1 to SD2 |

Inter-beat interval: time interval between successive R peaks in the ECG waveform.
NN interval: interval between two normal-to-normal inter-beat intervals.

#### Data pre-processing

The various HRV features have different ranges and orders of magnitude, which can have adverse affects on our modeling. To mitigate this, we standardized all features by removing the means and scaling to unit variance. We also applied a Yeo-Johnson power transformation to the features to minimize the effect of outliers on the learned model, and also improve the spread of the features.

### 3.5 Workload Classification

We first investigated whether UST HRV features were able to classify the subjective measure of workload. In total we had 12 datasets based on the interface and subject trial combinations. We created binary TLX labels for all our UST HRV data by mapping the weighted TLX scores to a label of *low* or *high* using the 50^th^ percentile as the threshold. The training dataset consisted of 14 UST HRV features and the binarized TLX label. Support vector classification with a linear kernel was used to predict the TLX label from the UST HRV features.

#### Feature Importance

We used feature ranking with recursive feature elimination and cross-validated selection of the best number of features to judge the importance of the various HRV features in terms of prediction accuracy. This approach was chosen to consider all features at once and also capture interactions between the features. Each classifier was fitted to the training datasets to yield separate rankings for the 14 UST HRV metrics. Stratified shuffle splitting was used to separate the dataset into training and validation sets in a 70-30 ratio with 10 fold cross-validation. The identified features were used in the final classification model.

### 3.6 HRV Trends For Different Trial Conditions

We also investigated how the various UST HRV features differed across trials with different conditions, including the baseline resting periods before and after the navigation tasks. Each trial was classified as *unsafe* if there were any collisions during task execution. For trials that did not have collisions, if the total trial time was greater than the median task time over all subjects, that trial was classified as *low performance*, and if the total trial time was less than the median, it was classified as *high performance*. With these classifications—unsafe, low performance, high performance, baseline (pre), and resting (post)—we performed a non-parametric Kruskal-Wallis test and the Conover’s post-hoc pairwise comparisons to find the strength of significance of any differences.

## 4 RESULTS AND DISCUSSION

We begin this section by presenting the results of the workload prediction accuracies. Next, we provide the statistical analysis results of the UST HRV features across different trial conditions for a select number of notable features. We used the non-parametric Kruskal-Wallis test to establish significance followed by Conover’s post-hoc pairwise comparisons if any significance was detected. For all figures, the notation * implies a p-value of *p* < 0.05, ** implies *p* < 0.01, and *** implies p < 0.001. Lastly, we discuss of our findings.

### 4.1 Prediction of TLX Workload

We performed the feature selection and classification on different datasets based on different combinations of the participant group and interface over both 30 second and 60 second windows, as shown in Figure 5. Although each dataset produced a different set of important features, *NNmode, NNvariance, NNmea, lf, SD*2, and *SampEn* were selected most frequently by the feature selection algorithm across all examined datasets.

**Figure 5:**
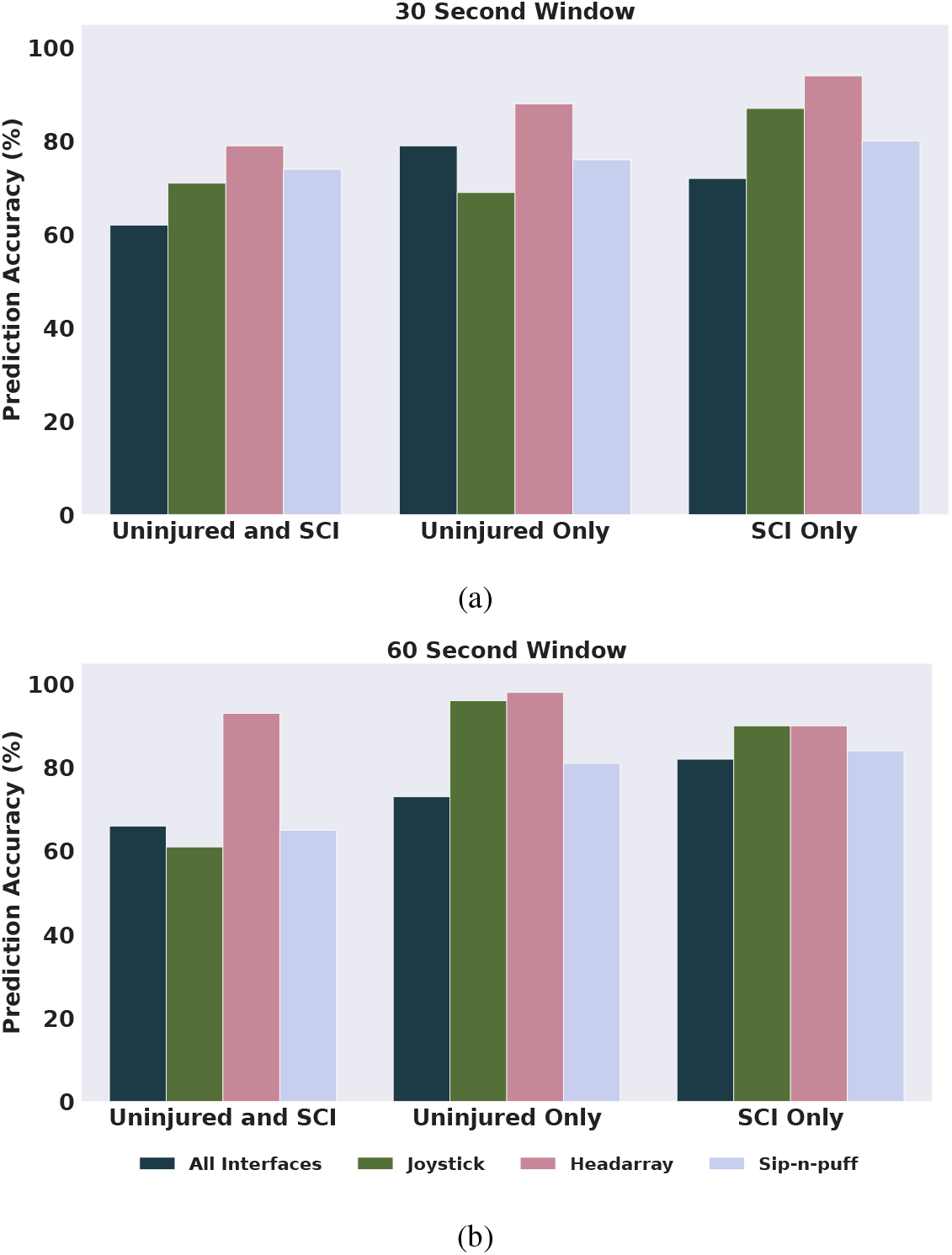
TLX prediction accuracy by participant demographic and interface. The grouping included combining SCI and uninjured participant data together, as well as looking at each group individually, and combining all three interfaces together as well as each interface individually. The classification was performed with UST HRV features measured over (a) 30s windows and (b) 60s windows.

Overall, prediction accuracies tend to be higher in datasets that included only one interface and one subject group, indicating differences in HRV features across interfaces that negatively affect the classifiers prediction power when grouped together. When comparing the prediction performance of UST HRV features measured over 30 versus 60 second windows, there was no significant difference with the latter being slightly more accurate. This is a promising finding, as the duration for many common wheelchair navigation tasks are less than one minute. Therefore, being able to predict workload over smaller time-windows is important for the utility of online workload prediction.

As seen in Figure 5, the classifier is consistently the most accurate in predicting the TLX workload score for trials using the headarray interface, across all participant groupings.

The trials with the joystick interface tended to be the fastest, so there were fewer windows over which UST HRV features could be calculated. Fewer windows resulted in sparser datasets for this interface, which may be one reason to explain the less accurate predictions compared to the headarray interface. Furthermore, the joystick interface requires less physical exertion compared the headarray, which may be another reason for the reduced prediction accuracy of workload using the HRV features.

With the sip-n-puff interface, the trial times were longest compared to the other interfaces, resulting in more windows per trial. From the perspective of the machine learning algorithm, this is a strength. However, the operating mechanism of the sip-n-puff interface—which requires inhaling and exhaling through a straw to provide commands—could affect the natural heart-rate rhythm which, in turn, can negatively affect the overall measured UST HRV features’ ability to predict workload. Exploratory scatter plots provided in the Appendix illustrate that the sip-n-puff data had very little-variance between different trials; evidence that this interface may not be well suited to using UST HRV features for the purpose of predicting workload.

### 4.2 Differences in HRV Trends

HRV signals were measured during five different experimental conditions, as described in Section III-F, for the same 12 datasets investigated in Section 4.1. Datasets that included the joystick and sip-n-puff interface showed no statistical significance for any UST HRV features between the five trial conditions.

We do observe statistical significance between various HRV features for trials with the headarray interface. A subset of these results are shown in Figure 6. For *RMSSD* and *SD*2, there was a significant difference during unsafe trials with collisions and the baseline resting state prior to the tasks. This suggests that UST HRV is correlated with unsafe conditions. There was also a significant difference in *lfhf* between unsafe and low performance driving conditions.

**Figure 6:**
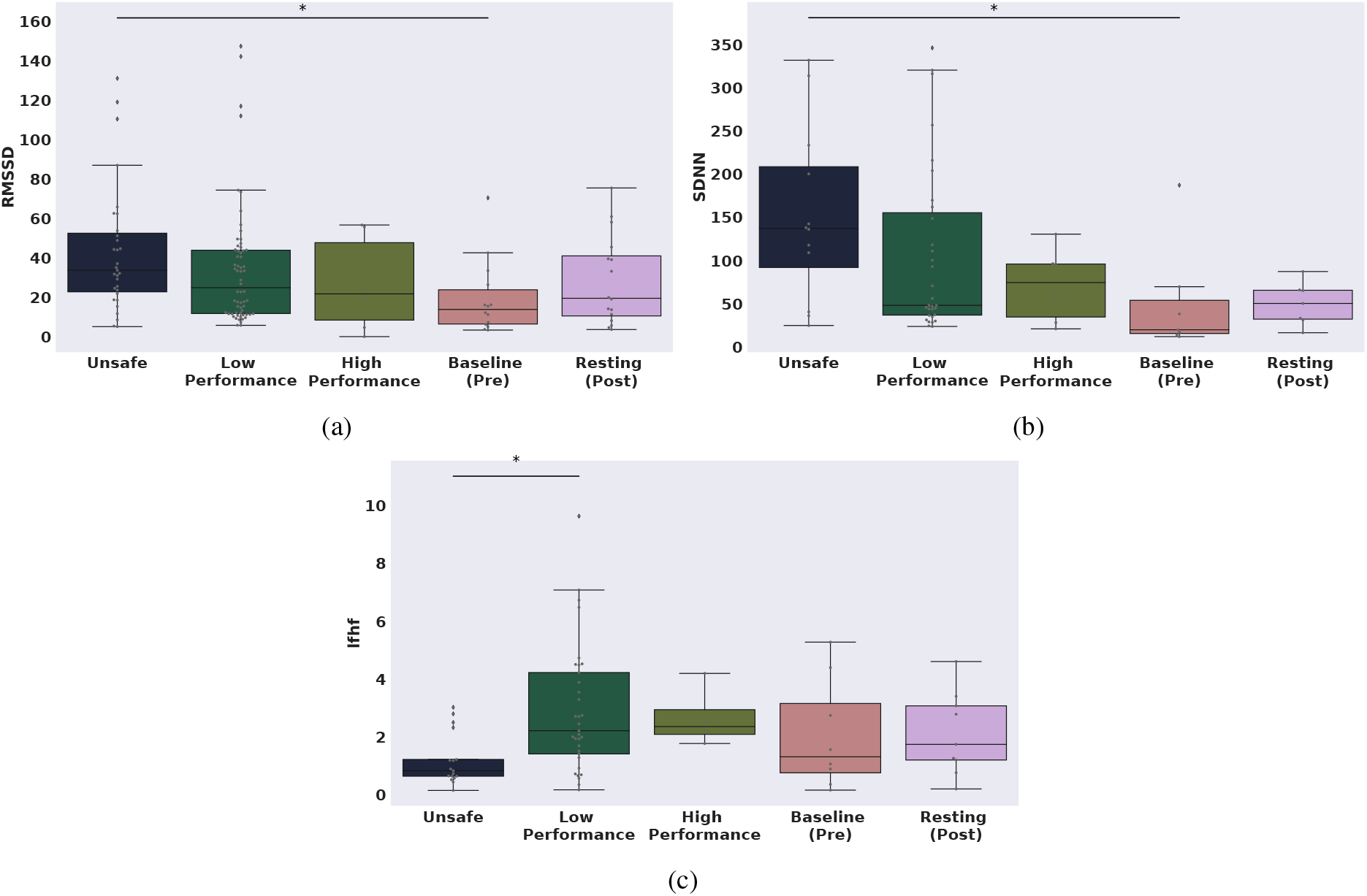
Difference in (a) RMSSD, (b) SDNN, and (c) lfhf measured over windows of 60 seconds using the headarray interface. There is a significant difference in these measures during baseline resting prior to the start of the experiment and during trials with collisions. Box plot showing the quartiles and distribution of the data, overlaid with scatter plot of the data. The notation * implies a p-value of *p* < 0.05, ** implies *p* < 0.01, and *** implies *p* < 0.001.

### 4.3 Discussion

Our results unveil a marked relationship between interface type and the density and spread of HRV features computed while a given interface is in use. The latter directly impacts the ability of a classifier to accurately predict workload scores using HRV features.

Our results further indicate that using any single UST HRV feature on its own may not be sufficient in predicting workload during wheelchair driving tasks. However, we are able to predict the workload with a model learned from a combination of ultra-short-term HRV features, with accuracies of over 90% for the interface associated with the greatest spread and density of HRV features (the headarray).

Our future work will investigate whether there is a causal relationship between changes in user input and changes in HRV. Also of interest is to investigate whether workload and changes in HRV have a causal or confounding relationship.

## 5 CONCLUSIONS

This study investigated the differences in HRV metrics during powered wheelchair navigation. To our knowledge, this was the first study that investigated the difference in UST HRV metrics during power wheelchair navigation by persons with and without spinal cord injury, using three commercially available interfaces that are each operated by different parts of body. We found evidence that the interface affects the HRV values and the ability to predict workload. Further studies are required to investigate the physiological phenomenon that affects how the physical operating mechanism of the interfaces affects the HRV and workload calculation, and whether the identified features of user input are reliable measure of workload.

## ACKNOWLEDGMENT

This material is based upon work supported by the National Science Foundation under Grant IIS-1552706. Any opinions, findings, and conclusions or recommendations expressed in this material are those of the authors and do not necessarily reflect the views of the National Science Foundation.

## APPENDIX

We include additional plots from the data analysis to support the discussion of our results.

As seen in Figure 7, exploratory scatter plots showed that the joystick data is sparse in comparison to the headarray data, which is quite dense. The joystick and headarray data both display a good amount of spread. The density of the data and the spread may explain why predictions using the headarray interface trials were the most accurate. On the other hand, the sip-n-puff data does not display much spread, which may explain why the classifiers suffered when using data collected from sip-n-puff trials.

**Figure 7:**
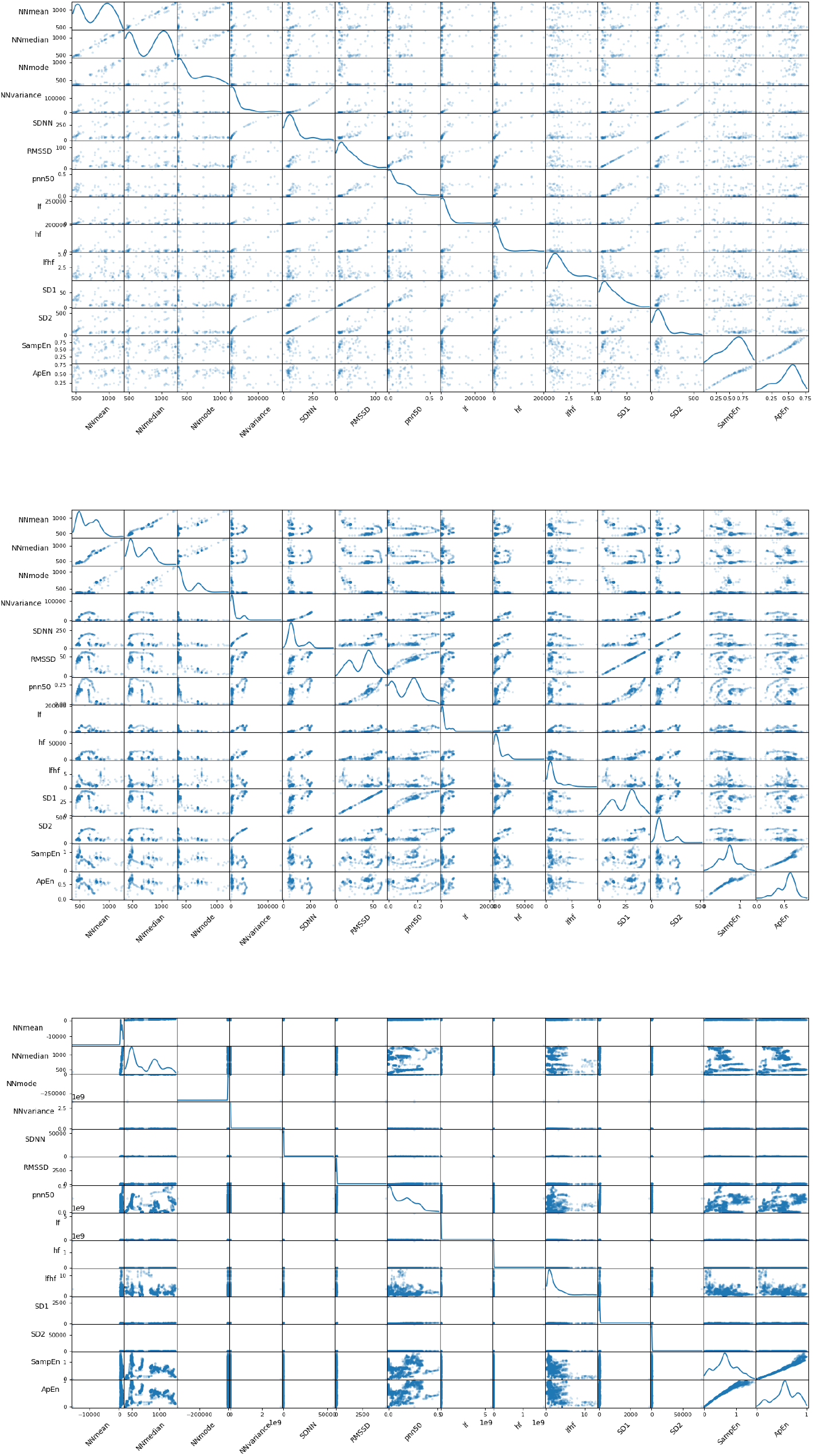
Scatter plots of all HRV features for joystick (top), headarray (middle), and sip-n-puff (bottom) interfaces. The headarray data is most dense and well spread. The joystick data is sparse and there is very little spread in the sip-n-puff data.

## References

[1] Mahdieh Nejati Javaremi, Michael Young, and Brenna Argall. Interface operation and implications for shared-control assistive robots. International Conference on Rehabilitation Robotics (ICORR), 2019.

[2] R. L. Charles and J. Nixon. Measuring mental workload using physiological measures: a systematic review. Applied Ergonomics, 2019.

[3] R. Castaldo, L. Montesinos, P. Melillo, C. James, and L Pecchia. Ultra-short term hrv features as surrogates of short term hrv: A case study on mental stress detection in real life. BMC Medical Informatics and Decision Making, 2019.

[4] S. Massaro and L. Pecchia. Heart Rate Variability (HRV) Analysis: A Methodology for Organizational Neuroscience. Organizational Research Methods, pages 354–393, 2019.

[5] D. Tao, H. Tan, H. Wang, X. Zhang, X. Qu, and T. Zhang. A systematic review of physiological measures of mental workload. International Journal of Environmental Research and Public Health, 2019.

[6] M. Soria-Oliver, J. S. Loṕez, and F. Torrano. Relations between mental workload and decision-making in an organizational setting. Psicologia: Reflexão e Crítica, 2018.

[7] G. Giannakakis, K. Marias, and M. Tsiknaisk. A stress recognition system using hrv parameters and machine learning techniques. International Conference on Affective Computing and Intelligent Interaction Workshops and Demos, pages 269–272, 2019.

[8] S. Delliaux, A. Delaforge, J. C. Deharo, and G. Chaumet. Mental workload alters heart rate variability, lowering non-linear dynamics. Frontiers in Physiology, 2019.

[9] R. McCraty and F. Shaffer. Heart rate variability: new perspectives on physiological mechanisms, assessment of self-regulatory capacity, and health risk. Global Advances in Health and Medicine, pages 46—–61, 2014.

[10] F. Shaffer and J. P. Ginsberg. An overview of heart rate variability metrics and norms. Frontiers in Public Health, 2017.

[11] T. Carlson and Y. Demiris. Collaborative control for a robotic wheelchair: Evaluation of performance, attention, and workload. IEEE Transactions on Systems, Man, and Cybernetics, Part B, pages 876–888, 2012.

[12] M.M. Lamti, HBen Khelifa and V. Hugel. Mental fatigue level detection based on event related and visual evoked potentials features fusion in virtual indoor environment. Cognitive Neurodynamics, pages 271–285, 2019.

[13] L Fehr, W E Langbein, and S B Skaar. Adequacy of power wheelchair control interfaces for persons with severe disabilities: a clinical survey. Journal of rehabilitation research and development, pages 353–60, 2000.

[14] S. G. Hart. Nasa-task load index (nasa-tlx); 20 years later. Proc. of the Human Factors and Ergonomics Society Annual Meeting, pages 904–908, 1985.

[15] J. Pan and W. J. Tompkins. A real-time qrs detection algorithm. IEEE Transactions on Biomedical Engineering, pages 230–236, 1985.

[16] A. Vest, G. Da Poian, Q. Li, C. Liu, S. Nemati, A. Shah, and G. D. Clifford. An open source benchmarked toolbox for cardiovascular waveform and interval analysis. Physiological Measurement, 2018.

[17] F. Shaffer, Z. Meehan, and C Zerr. A critical review of ultra-short-term heart rate variability norms research. Frontiers in Neuroscience, pages 1–11, 2020.

[18] M. R. Esco and A. A. Flatt. Ultra-short-term heart rate variability indexes at rest and post-exercise in athletes: Evaluating the agreement with accepted recommendations. Journal of Sports Science Medicine, pages 535–541, 2014.

[19] L. Salahuddin, J. Cho, M. G. Jeong, and D. Kim. Ultra short term analysis of heart rate variability for monitoring mental stress in mobile setting. IEEE Engineering in Medicine and Biology Society, pages 4656–4659, 2014.

[20] M. Munoz, A. Van Roon, H. Riese, C. Thio, E. Oostenbroek, I. Westrik, E. De Geus, R. Gansevoort, J. Lefrandt, I. Nolte, and H. Snieder. Validity of (ultra-)short recordings for heart rate variability measurements. PLoS ONE, pages 1–15, 2015.

